# Transethnic genetic correlation estimates from summary statistics

**DOI:** 10.1101/036657

**Authors:** Brielin C. Brown, Asian Genetic Epidemiology Network-Type 2 Diabetes (AGEN-T2G) Consortium, Chun Jimmie Ye, Alkes L. Price, Noah Zaitlen

## Abstract

The increasing number of genetic association studies conducted in multiple populations provides unprecedented opportunity to study how the genetic architecture of complex phenotypes varies between populations, a problem important for both medical and population genetics. Here we develop a method for estimating the *transethnic genetic correlation*: the correlation of causal variant effect sizes at SNPs common in populations. We take advantage of the entire spectrum of SNP associations and use only summary-level GWAS data. This avoids the computational costs and privacy concerns associated with genotype-level information while remaining scalable to hundreds of thousands of individuals and millions of SNPs. We apply our method to gene expression, rheumatoid arthritis, and type-two diabetes data and overwhelmingly find that the genetic correlation is significantly less than 1. Our method is implemented in a python package called *popcorn*.

## Introduction

Many complex human phenotypes vary dramatically in their distributions between populations due to a combination of genetic and environmental differences. For example, northern Europeans are on average taller than southern Europeans^1^ and African Americans have an increased rate of hypertension relative to European Americans^2^. The genetic contribution to population phenotypic differentiation is driven by differences in causal allele frequencies, effect sizes, and genetic architectures. Understanding the root causes of phenotypic differences worldwide has profound implications for biomedical and clinical practice in diverse populations, the transferability of epidemiological results, aiding multi-ethnic disease mapping^3,4^, assessing the contribution of non-additive and rare variant effects, and modeling the genetic architecture of complex traits. In this work we consider a central question in the global study of phenotype: do genetic variants have the same phenotypic effects in different populations?

While the vast majority of GWAS have been conducted in European populations^5^, the growing number of non-European and multi-ethnic studies^4,6,7^ provide an opportunity to study genetic effect distributions across populations. For example, one recent study used mixed-model based methods to show that the genome-wide genetic correlation of schizophrenia between European and African Americans is nonzero^8^. While powerful, computational costs and privacy concerns limit the utility of genotype-based methods. In this work, we make two significant contributions to studies of transethnic genetic correlation. First, we expand the definition of genetic correlation to better account for a transethnic context. Second, we develop an approach to estimating genetic correlation across populations that uses only summary level GWAS data. Similar to other recent summary statistics based methods^99–20^, our approach supplements summary association data with linkage disequilibrium (LD) information from external reference panels, avoids privacy concerns, and is scalable to hundreds of thousands of individuals and millions of markers. Unlike traditional approaches that focus on the similarity of GWAS results^21–25^ we utilize the entire spectrum of GWAS associations while accounting for LD in order to avoid filtering correlated SNPs.

In a single population, the genetic correlation of two phenotypes is defined as the correlation coefficient of SNP effect sizes^18,26^. In multiple populations, differences in allele frequency motivate multiple possible definitions of genetic correlation. Because a variant may have a higher effect size but lower frequency in one population, we consider both the correlation of allele effect sizes as well as the correlation of allelic impact. We define the *transethnic genetic effect correlation* (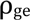, previously defined by Lee et al^26^ and implemented in GCTA) as the correlation coefficient of the per-allele SNP effect sizes, and the *transethnic genetic impact correlation* (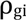) as the correlation coefficient of the population-specific allele variance normalized SNP effect sizes.

Intuitively, the genetic effect correlation measures the extent to which the same variant has the same phenotypic change, while the genetic impact correlation gives more weight to common alleles than rare ones separately in each population. Consider the case of a SNP that is rare in population 1 but common in population 2, and has an identical effect size in both populations. In this case, the correlation of effect sizes (the genetic effect correlation 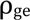) is 1, but this provides an incomplete picture of the relationship between the two populations, as the allele has a much bigger impact on the distribution of the phenotype in population 2. Therefore, we define the genetic impact correlation 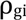 as the correlation of effect sizes after normalizing genotypes to have mean 0 and variance 1. In our hypothetical case 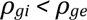, however the opposite can also be true. Consider again the case of a SNP rare in population 1 but common in population 2. If the effect size is large in the first population but small in the second, then 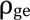 may be much less than 1, but the impact of the allele in the two populations will be similar and therefore 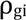 will be close to 1. While other definitions of the genetic correlation are possible (see discussion), these quantities capture two important questions about the study of disease in multiple populations: to what extent do the same mutations in multiple populations differ in their phenotypic effects? And, to what extent are these differences mitigated or exacerbated by differences in allele frequency?

To estimate genetic correlation, we take a Bayesian approach wherein we assume genotypes are drawn separately from within each population and effect sizes have a normal prior (the infinitesimal model^27^). While unlikely to represent reality, this model has been used successfully in practice^8,16,17,28,29^. The infinitesimal assumption yields a multivariate normal distribution on the observed test statistics (Z-scores), which is a function of the heritability and genetic correlation. Rather than pruning SNPs in LD^10,30,31^, this allows us to explicitly model the resulting inflation of Z-scores. We then maximize an approximate weighted likelihood function to find the heritability and genetic correlation. This method is implemented in a python package called *popcorn*. Though derived for quantitative phenotypes, *popcorn* extends easily to binary phenotypes under the liability threshold model. We show via extensive simulation that *popcorn* produces unbiased estimates of the genetic correlation and the population specific heritabilities, with a standard error that decreases as the number of SNPs and individuals in the studies increases. Furthermore, we show that our approach is robust to violations of the infinitesimal assumption.

We apply *popcorn* to European and Yoruban gene expression data^32^ as well as GWAS summary statistics from European and East Asian rheumatoid arthritis and type-two diabetes cohorts ^33,34^. Our analysis of gEUVADIS shows that our summary statistic based estimator is concordant with the mixed model based estimator. We find that the mean transethnic genetic correlation across all genes is low 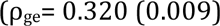, but increases substantially when the gene is highly heritable in both populations 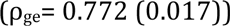. in RA and T2D, we find the genetic effect correlation to be 0.463 (0.058) and 0.621 (0.088), respectively.

Across all phenotypes considered, we overwhelmingly find that the transethnic genetic correlation is significantly less than one. This observation highlights the need to study phenotypes in multiple populations as it implies that, up to the effects of un-observed variants, effect sizes at common SNPs tend to differ between populations. This indicates that GWAS results may not transfer between populations, and therefore disease risk prediction in non-Europeans based on current GWAS results may be problematic, necessitating a multi-population approach to gain insight into inter-population differences in the genetic architecture of complex traits.

## Methods

Our method takes as input summary association statistics from two studies of a phenotype in two different populations, along with two sets of reference genotypes each matching one of the populations in the study. Our method has two steps: first, we estimate the diagonal elements of the LD matrix products 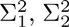 and 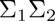 then, using these estimates, we find the maximum likelihood values and estimate standard errors of the parameters of interest: 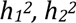 and 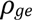 or 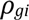. The details follow.

Consider GWAS of a phenotype conducted in two different populations. Assume we have *N*_1_ individuals genotyped on *M* SNPs in study one and *N_2_* individuals genotyped on the same SNPs study two. Let *X_1_*, *X_2_* be the matrices of mean-centered genotypes in study one and study two, respectively, and let *Y_1_*, *Y_2_* be their normalized phenotypes. Let *f_1_*, *f_2_* be vectors of the allele frequencies of the *M* SNPs common to both populations. Assuming Hardy-Weinberg equilibrium within each population separately, the allele variances are 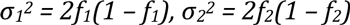. Let *β_1_*, *β_2_* be the (unobserved) per-allele effect sizes for each SNP in studies one and two, respectively. The heritability in study one is then 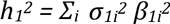 (and likewise for study two). The objective of this work is to estimate transethnic genetic correlation from summary statistics of common variants 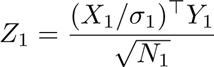 (and likewise for study two) and estimates of population LD matrices (*Σ_1_* and *Σ_2_*) from external reference panels. Define the *genetic effect correlation* 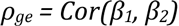 and the *genetic impact correlation* 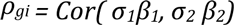.

We assume the genotypes are drawn randomly from each population and that phenotypes are generated by the linear model 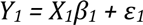 (likewise for phenotype two). When effect sizes *β* are assumed inversely proportional to allele frequency, as is commonly done^16,29^, we show (Appendix) that under the linear infinitesimal genetic architecture, the joint distribution of the Z-scores from each study is asymptotically multivariate normal with mean 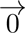 and variance:

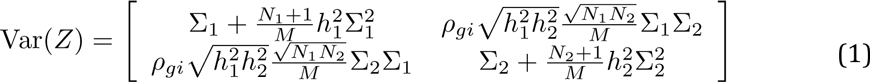

However, when effect sizes are assumed independent of allele frequency we show:

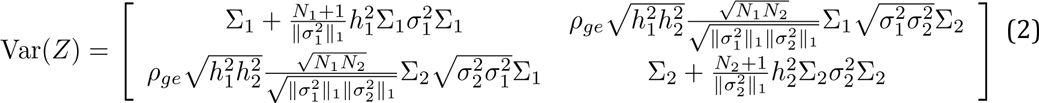

Given these equations for variance, the quantities 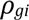 or 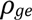 and 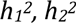 can be estimated by maximizing the multivariate normal likelihood, 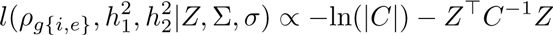 is either of the above covariance matrices (1) and (2). Because *Σ_1_* and *Σ_2_* are estimated from finite external reference panels, maximum likelihood estimation of the above multivariate normal leads to over-fitting. We employ two optimizations to avoid this problem. First, we maximize an approximate weighted likelihood that uses only the diagonal elements of each block of *Var(Z)*. This allows us to account for the LD-induced inflation of tests statistics, but discards covariance information between pairs of Z-scores, and therefore leads to over-counting Z-scores of SNPs in high LD. To compensate for this, we down weight Z-scores of SNPs in proportion their LD. Second, rather than compute the full products 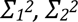 and 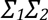 over all *M* SNPs in the genome, we choose a window size *W* and approximate the product by 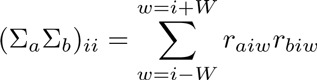. These optimizations are similar to those employed by LD score regression^16^. The full details of the derivation and optimization are provided in the appendix.

## Results

### Simulated Genotypes and Simulated Phenotypes

We simulated 50,000 European-like (EUR) and 50,000 East Asian-like (EAS) individuals at 248,953 SNPs from chromosomes 1-3 with allele frequency above 1% in both European and East Asian HapMap3 populations with HapGen2^35^. HapGen2 implements a haplotype recombination with mutation model that results in excess local relatedness among the simulated individuals. To account for this local structure, we used Plink2^36^ to filter individuals with genetic relatedness above 0.05, resulting in 4499 EUR-like indivudals and 4837 EAS-like individuals. From these simulated individuals, 500 per population were chosen uniformly at random to serve as an external reference panel for estimating *Σ_1_* and *Σ_2_*.

In each simulation effect sizes were drawn from a “spike and slab” model, where 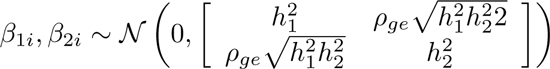 with probability *p* and 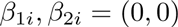 with probability *1*-*p*.*ρ_gi_* was analytically computed from the simulated effect sizes and allele frequencies in the simulated reference genotypes. Quantitative phenotypes were generated under a linear model with i.i.d. noise and normalized to have mean 0 and variance 1, while binary phenotypes were generated under a liability threshold model where individuals are labeled cases when their liability exceeds a threshold 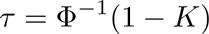, with K the population disease prevalence^37^.

We varied 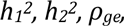 and 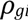, as well as the number of individuals in each study (*N_1_*, *N_2_*), the number of SNPs (*M*), the population prevalence *K*, and proportion of causal variants (*p*) in the simulated GWAS and generated summary statistics for each study. The results shown in Figure 1 and Figure S1 demonstrate that the estimators are nearly unbiased as the genetic correlation and heritabilities vary. Furthermore, by varying the proportion of causal variants *p* we show that our estimator is robust to violations of the infinitesimal assumption (Figure S2). In figure S3, we show that the standard error of the estimator decreases as the number of SNPs and individuals in the study increases. Finally, we show in Table S1 that our estimates of the heritability of liability in case control studies are nearly unbiased.

**Figure 1:**
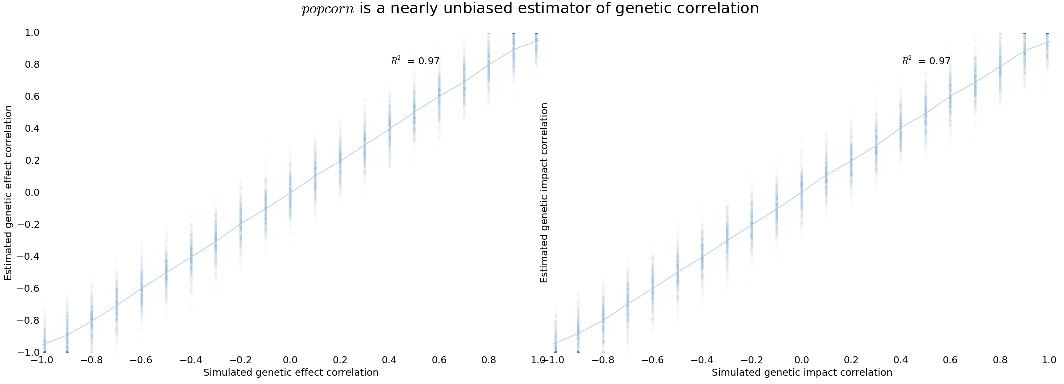
True and estimated genetic impact and effect correlation. All simulations conducted with simulated EUR and EAS heritability of 0.5 using 4499 simulated EUR and 4836 simulated EAS individuals at 248,953 SNPs.

### Simulations with nonstandard disease models

Our approach, as well as genotype-based methods such as GCTA, makes assumptions about the genetic architecture of complex traits. Previous work has shown that violations of these assumptions can lead to bias in heritability estimation^38^, therefore we sought to quantify the extent that this bias may effect our estimates. We simulated phenotypes under six different disease models. *Independent*: effect size independent of allele frequency. *Inverse*: effect size inversely proportional to allele frequency. *Rare*: only SNPs with allele frequency under 10% affect the trait. *Common*: only SNPs with allele frequency between 40% and 50% affect the trait. *Difference*: effect size proportional to difference in allele frequency. *Adversarial*: difference model with sign of beta set to increase the phenotype in the population where the allele is most common. Additional genetic architectures are possible, including ones where effect sizes are not a direct function of MAF^39^.

We simulated phenotypes using genotypes with allele frequency above 1% or 5% and compared the true and estimated genetic impact and effect correlation among all models (Table 1). We find that when only SNPs with frequency above 5% in both populations are used, the difference in 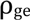 and 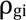 is minimal except in the most adversarial cases. Even in the adversarial model, the true difference is only 7%. Though unlikely to represent reality, the four nonstandard disease models result in substantial bias in our estimators. When SNPs with allele frequency above 1% in both populations are included, the differences are more pronounced. This is because the normalizing constant 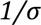 rapidly increases as the SNP becomes more rare. Indeed, as SNPs become more rare having an accurate disease model becomes increasingly important. Therefore we proceed with a 5% MAF cutoff in our analysis of real data, and use the notation 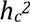 to refer to the heritability of SNPs with allele frequency above 5% in both populations (the *common-SNP heritability*). Note, however, that one of the advantages of maximum likelihood estimation in general is that the likelihood can be reformulated to mimic the disease model of interest.

**Table 1:** True and estimated values of the genetic impact and effect correlation in simulated EUR-like and EAS-like genotypes. Results are the average of 100 simulations with phenotype heritability of 0.5 in each population.

|  | MAF > 0.01 |  |  |  | MAF > 0.05 |  |  |  |
| --- | --- | --- | --- | --- | --- | --- | --- | --- |
| Model | $\rho_{ge}$ | $\rho_{gi}$ | $\hat{\rho}_{ge}$ | $\hat{\rho}_{gi}$ | $\rho_{ge}$ | $\rho_{gi}$ | $\hat{\rho}_{ge}$ | $\hat{\rho}_{gi}$ |
| Independent | 0.500 | 0.478 | 0.500 | 0.460 | 0.500 | 0.488 | 0.509 | 0.469 |
| Inverse | 0.431 | 0.500 | 0.567 | 0.496 | 0.479 | 0.500 | 0.555 | 0.482 |
| Rare | 0.500 | 0.467 | 0.382 | 0.863 | 0.500 | 0.496 | 0.998 | 0.756 |
| Common | 0.500 | 0.500 | 0.522 | 0.493 | 0.500 | 0.500 | 0.502 | 0.496 |
| Difference | 0.500 | 0.416 | 0.354 | 0.435 | 0.500 | 0.461 | 0.410 | 0.412 |
| Adversarial | 0.710 | 0.604 | 0.525 | 0.651 | 0.714 | 0.667 | 0.601 | 0.675 |

### Validation of Popcorn using gene expression in GEUVADIS

We compared the common-SNP heritability 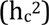 and genetic correlation estimates of *popcorn* to GCTA in the gEUVADIS dataset for which raw genotypes are publicly available. gEUVADIS consists of RNA-seq data for 464 lymphoblastoid cell line (LCL) samples from five populations in the 1000 genomes project. Of these, 375 are of European ancestry (CEU, FIN, GBR, TSI) and 89 are of African ancestry (YRI). Raw RNA-sequencing reads obtained from the European Nucleotide Archive were aligned to the transcriptome using UCSC annotations matching hg19 coordinates. RSEM was used to estimate the abundances of each annotated isoform and total gene abundance is calculated as the sum of all isoform abundances normalized to one million total counts or transcripts per million (TPM). For eQTL mapping, Caucasian and Yoruban samples were analyzed separately. For each population, TPMs were median-normalized to account for differences in sequencing depth in each sample and standardized to mean 0 and variance 1. Of the 29763 total genes, 9350 with TPM > 2 in both populations were chosen for this analysis.

For each gene, we conducted a *cis*-eQTL association study at all SNPs within 1 megabase of the gene body with allele frequency above 5% in both populations using 30 principal components as covariates. We found that GCTA and *popcorn* agree on the global distribution of heritability (Figure S3), and that GCTA’s estimates of genetic correlation have a similar distribution to *popcorn*’s genetic effect and genetic impact correlation estimates (Figure 2). While the number of SNPs and individuals included in each gene analysis are too small to obtain accurate point estimates of the genetic correlation on a pergene basis (N=464, M=4279.5), the large number of genes enables accurate estimation of the global mean heritability and genetic correlation.

**Figure 2:**
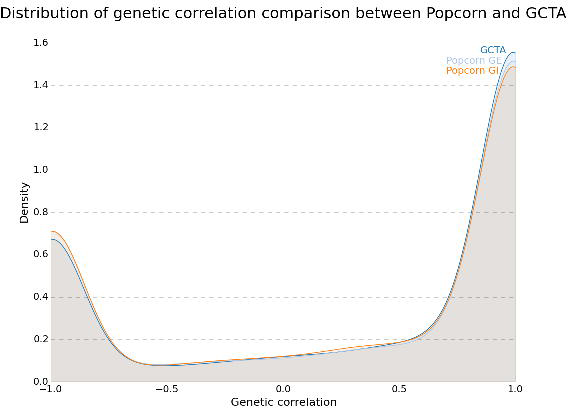
Distribution of genetic correlation comparison between popcorn and GCTA. Distribution was computed using a gaussian kde on the set of genetic correlation estimates.

### Common-SNP heritability and genetic correlation of gene expression in gEUVADIS

We find that the average 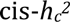 of the expression of the genes we analyzed was 0.093 (0.002) in EUR and 0.088 (0.002) in YRI. Our estimates are higher than previously reported average cis-heritability estimates of 0.055 in whole blood and 0.057 in adipose^40^, which could arise for several reasons. First, we remove 68% of the transcripts that are lowly expressed in either the YRI or EUR data. Second, estimates from RNA-seq analysis of cell lines might not be directly comparable to microarray data from tissue.

The average genetic effect correlation was 0.320 (0.010), while the average genetic impact correlation was 0.313 (0.010). Notably, the genetic correlation increases as the 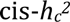 of expression in both populations increases (Figure 3). In particular, when the 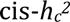 of the gene is at least 0.2 in both populations the genetic effect correlation was 0.772 (0.017) while the genetic impact correlation was 0.753 (0.018).

**Figure 3:**
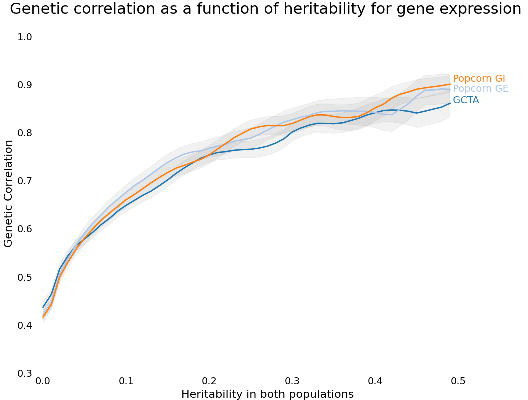
Genetic correlation as a function of heritability for gene expression. The mean and standard error of the genetic correlation of the set of genes with 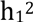 and 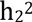 exceeding threshold *h* in each analysis (y-axis] is plotted against *h* (x-axis].

In order to verify that there were no small-sample size or conditioning biases in our analysis, we analyzed the genetic correlation of simulated phenotypes over the gEUVADIS genotypes. We sampled pairs of heritabilities from the estimated expression heritability distribution and simulated pairs of phenotypes to have the given heritability and a genetic effect correlation of 0.0 over randomly chosen 4000 base regions from chromosome 1 of the gEUVADIS genotypes. Without conditioning, the average estimated genetic effect correlation was -0.002 (0.003), indicating that the estimator remained unbiased. Furthermore, the average estimated genetic effect correlation was not significantly different from 0.0 conditional on the estimates of heritability being above a certain threshold (Figure S4).

We find that while the average genetic correlation is low, the genetic correlation increases with the 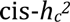 of the gene, indicating that as cis-genetic regulation of gene expression increases it does so similarly in both YRI and EUR populations. This may help interpret the recent observation that while the global genetic correlation of gene expression across tissues is low^40^, *cis*-eQTL’s tend to replicate across tissues^41^. As the presence of a *cis*-eQTL indicates substantial cis-genetic regulation, an analysis of eQTL replication across tissues is implicitly conditioning on the heritability of gene expression being high and therefore may indicate much higher genetic correlation than average.

### Summary statistics of RA and T2D

Finally, we sought to examine the transethnic 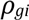 and 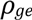 in RA and T2D cohorts for which raw genotypes are not available. We obtained summary statistics of GWAS for rheumatoid arthritis and type-2 diabetes conducted in European and East Asian populations. We used genotypes from 504 East Asian and 503 European individuals sequenced as part of the 1000 genomes project as population-specific external reference panels for our EAS and EUR summary statistics, respectively. We removed the MHC region (chromosome 6, 25-35 Mb) from the RA summary statistics. We estimated the common-SNP heritability and genetic correlation using 2,539,629 SNPs genotyped or imputed in both RA studies and 1,054,079 SNPs genotyped or imputed in both T2D studies with allele frequency above 5% in 1000 genomes EUR and EAS populations. The 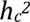 and genetic correlation estimates are presented in Table 2. Our RA 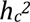 estimates of 0.177 (0.015) and 0.221 (0.026) for EUR and EAS, respectively, are lower than a previously reported mixed-model based heritability estimates of 0.32 (0.037) in Europeans^42^. Similarly, our T2D 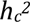 estimates of 0.242 (0.013) and 0.105 (0.021) for EUR and EAS, respectively, are lower than a previously reported mixed-model based estimate of 0.51 (0.065) in Europeans^42^. We stress that this discrepancy is likely due to the difference between common-SNP heritability 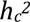 and total narrow-sense heritability 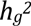. Furthermore, estimates of the heritability of T2D from family studies can vary significantly^43,44^.

**Table 2:** Heritability and genetic correlation of RA and T2D between EUR and EAS populations. EUR RA data contained 8,875 cases and 29,367 controls for a study prevalence of 0.23. EAS RA data contained 4,873 cases and 17,642 controls for a study prevalence of 0.22. RA disease prevalence was assumed to be 0.5% in both populations^7^. T2D EUR data contained 12171 cases and 56862 controls for a study prevalence of 0.18. T2D EAS data contained 6952 cases and 11865 controls for a study prevalence of 0.37. T2D EUR prevalence was assumed to be 8%^33^ while T2D EAS prevalence was assumed to be 9%^48^.

| | | $h_{EUR}^2$ liability | $h_{EAS}^2$ liability | $\rho_{ge}$ | $\rho_{gi}$ |
| --- | --- | --- | --- | --- | --- |
| RA | Est. (SE) | 0.18 (0.02) | 0.22 (0.03) | 0.46 (0.06) | 0.46 (0.06) |
|  | 95% CI | [0.15, 0.21] | [0.16, 0.28] | [0.34, 0.58] | [0.34, 0.58] |
| | $p_{>0}$ | 3.90e-32 | 1.89e-17 | 1.37e-15 | 8.16e-16 |
| | $p_{<1}$ | 0.0 | 3.1e-197 | 2.53e-20 | 4.87e-22 |
| T2D | Est. (SE) | 0.24 (0.01) | 0.11 (0.02) | 0.62 (0.09) | 0.61 (0.08) |
|  | 95% CI | [0.22, 0.26] | [0.07, 0.15] | [0.44, 0.80] | [0.45, 0.77] |
| | $p_{>0}$ | 2.41e-77 | 5.73e-7 | 1.70e-12 | 2.85e-13 |
| | $p_{<1}$ | 0.0 | 0.0 | 1.066e-05 | 2.06e-06 |

We find the genetic effect correlation in RA and T2D to be 0.463 (0.058) and 0.621 (0.088), respectively, while the genetic impact correlation is not significantly different at 0.455 (0.056) and 0.606 (0.083). The transethnic genetic impact and effect correlation for both phenotypes are significantly different from both 1 and 0 (Table 2), showing that while there is clear genetic overlap between the phenotypes, the per allele effect sizes differ significantly between the two populations.

## Discussion

We have developed the transethnic genetic effect and genetic impact correlation and provided an estimator for these quantities based only on summary-level GWAS information and suitable reference panels. We have applied our estimator to several phenotypes: rheumatoid arthritis, type-2 diabetes and gene expression. While the gEUVADIS dataset lacks the power required to make inferences about the genetic correlation of single or small subsets of genes, we can make inferences about the global structure of genetic correlation of gene expression. We find that the global mean genetic correlation is low, but that it increases substantially when the heritability is high in both populations. In all phenotypes analyzed, the genetic correlation is significantly different from both 0 and 1. Our results show that global differences in SNP effect size of complex traits can be large. In contrast, effect sizes of gene expression appear to be more conserved where there is strong genetic regulation.

It is not possible to draw conclusions about polygenic selection from estimates of transethnic genetic correlation. The effect sizes may be identical 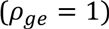 while polygenic selection acts to change only the allele frequencies. Similarly, the effect sizes may be different 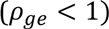 without selection. Differences in effect sizes at common SNPs can result from many phenomena. We expect un-typed and un-imputed variants differentially linked to observed SNPs to contribute significantly, along with rare or population-specific variants differentially linked to observed SNPs. If a gene-gene or gene-environment interaction exists, but only marginal effects are tested, the observed marginal effects may be different in each population due to allele frequency differences even if the interaction effect is the same in both populations, and this will result in decreased genetic correlation. While within-locus (dominance) interactions may also play a role^45^, the magnitude of this effect has been debated^46^. We emphasize that we cannot differentiate between these effects on the basis of this analysis alone, and further research is required to establish the magnitude of the contribution of each of these effects to inter-population effect size differences.

Estimates of the transethnic genetic correlation are important for several reasons. They may help inform best practices for transethnic meta-analysis, potentially offering improvements over current methods that use F_st_ to cluster populations for analysis^4^. Further, the transethnic genetic correlation constrains the limit of out of sample phenotype predictive power. If the maximum within population correlation of predicted phenotype *P* to true phenotype *Y* is 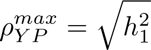, then the maximum out of population correlation is 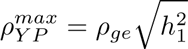 (Appendix). Our observation that for RA, T2D, and gene expression the genetic correlation is low shows that out of population phenotypic predictive power is quite low. Similarly, it implies that disease risk assessment in non-Europeans based on current GWAS results may be problematic, necessitating increased study of disease in many populations to gain insight into differences in genetic architecture and improve risk assessment.

While the genetic correlation of multiple phenotypes in one population has a relatively straightforward definition, extending this to multiple populations motivates multiple possible extensions. In this work we have provided estimators for the correlation of genetic effect and genetic impact but other quantities related to the shared genetics of complex traits between populations include the correlation of variance explained 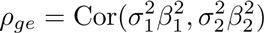 and proportion of shared causal variants between the two populations. Interestingly, while our goal was to construct an estimator that determined the extent of genetic sharing independent of allele frequency, we observe that the correlation of genetic effect and genetic impact are similar. Furthermore, our simulations show that under a random effects model utilizing only SNPs with allele frequency above 5% in both populations the true genetic effect and genetic impact correlation are similar. We conclude that at variants common in both populations, differences in effect size and not allele frequency are driving the transethnic phenotypic differences in these traits.

Our approach to estimating genetic correlation has two major advantages over mixed-model based approaches. First, utilizing summary statistics allows application of the method without data-sharing and privacy concerns that come with raw genotypes. Second, our approach is linear in the number of SNPs avoiding the computational bottleneck required to estimate the genetic relationship matrix. Conceptually, our approach is very similar to that taken by LD score regression. Indeed, the diagonal of the LD matrix product in one population are exactly the LD-scores 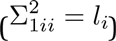. One could ignore our likelihood-based approach and define *cross-population scores* 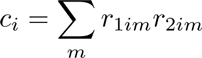 in order to exploit the linear relationship 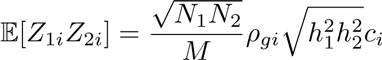 (a similar approach can be taken for the genetic effect correlation). Since LD-score regression has been successfully used to compute the genetic correlation of two phenotypes in a single population, this derivation can be viewed as an extension of LD-score regression to one phenotype in two different populations. The main difference in our approach is choosing maximum likelihood rather than regression in order to fit the model. A comparison of our method to the ldsc software shows they perform similarly as heritability estimators (Figure S5).

Of course, our method is not without drawbacks. First, it requires a large sample size and large number of SNPs to achieve standard errors low enough to generate accurate estimates. Until recently large sample GWAS have been rare in non-European populations, though they are becoming more common. Similarly, reference panel quality may suffer in non-European populations and this may impact downstream analysis^47^. Second, it is limited to analyzing relatively common SNPs, both because having an accurate disease model is important for the analysis of rare variants and because effect size and correlation coefficient estimates have a high standard error at rare SNPs^16^. Third, our analysis is currently limited to SNPs that are present in both populations. Indeed it is currently unclear how best to handle population-specific variants in this framework. Fourth, our estimator of *ρ* is bounded between -1 and 1. This may induce bias when the true value is close to the boundary and the sample size is small. Finally, admixed populations induce very long-range LD that is not accounted for in our approach and we are therefore limited to un-admixed populations^16^.

Our analysis leaves open several avenues for future work. We approximately maximize the likelihood of an *M*×*M* multivariate normal distribution via a method that uses only the diagonal elements of each block, discarding covariance information between Z-scores. A better approximation may lower the standard error of the estimator, facilitating an analysis of the genetic correlation of functional categories, pathways and genetic regions. We would also like to extend our analysis to include population specific variants as well as variants at frequencies between 1-5% or lower than 1%. Our simulations indicate that having an accurate disease model is important for determining the difference between the genetic effect and genetic impact correlation when rare variants are included. Maximum likelihood approaches are well suited to different genetic architectures. For example, one could estimate both the global relationship between allele frequency and effect size and the global relationship between per-SNP F_ST_ and genetic correlation by incorporating parameters α and γ into the prior distribution of the effect sizes, 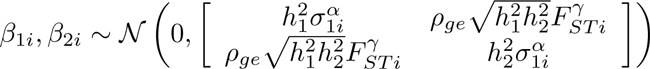. We expect that incorporating these parameters will improve estimates of heritability and genetic correlation while revealing important biological insights.

## Appendix

Consider two GWAS of a phenotype conducted in different populations populations. Assume we have *N_1_* individuals genotyped or imputed to *M* SNPs in study one and *N_2_* individuals genotyped or imputed to *M* SNPs in study two. Let *X_1_*, *X_2_* and *Y_1_*, *Y_2_* be the matrices of mean-centered genotypes and phenotypes of the individuals in study one and two, respectively, with *f*_1_, *f*_2_ the allele frequencies of the *M* SNPs common to both populations. Assuming Hardy-Weinberg equilibrium, the allele variances are 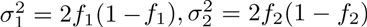. Let *β_1_*, *β_2_* be the (unobserved) per-allele effect sizes for each SNP in studies one and two, respectively. Define the *genetic impact correlation* 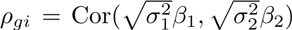 and the *genetic effect correlation* 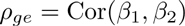. We present a maximum likelihood framework for estimating the heritability of the phenotype in study 1 and it’s standard error, the heritability of the phenotype in study 2 and it’s standard error, and the genetic effect and impact correlation of the phenotype between the studies and it’s standard error given only the summary statistics *Z_1_*, *Z_2_* and reference genotypes *G_1_*, *G_2_* representing the populations in the studies. We assume that genotypes are drawn randomly from populations with expected correlation matrices *Σ_1_* (and similarly for study two), and that every SNP is causal with a normally distributed effects size (though this assumption is not necessary in practice, see Figure S1).

### Genetic impact correlation

Let 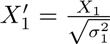 (and similarly for study 2) be normalized genotype matrices. We consider the standard linear model for generation of the phenotypes, where

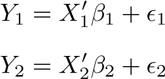

For convenience of notation let 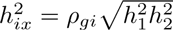. We assume the SNP effects follow the infinitesimal model, where every SNP has an efect size drawn from the normal distribution, and that the residuals are independent for each individual and normally distributed:

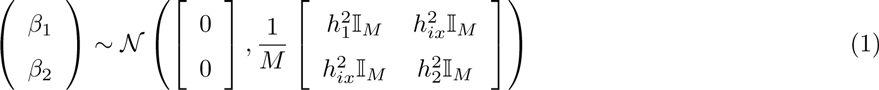

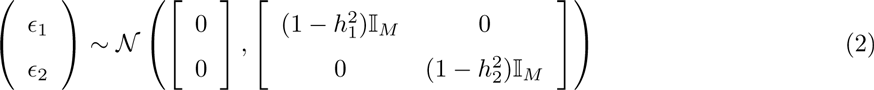

where 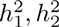 are the heritability of the disease in study one and two, respectively, and 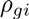 is the genetic impact correlation.

Using the above model, we compute the distribution of the observed *Z* scores as a function of the reference panel correlations and the model parameters 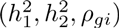. Given a distribution for *Z* and an observation of *Z* we can then choose parameters which give the highest probability of observing *Z*. First, we compute the distribution of *Z*. It is well known that the *Z*-scores of a linear regression are normally distributed given *β* when the sample size is large enough. Since 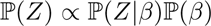 and the product of normal distributions is normal, we only need to compute the unconditional mean and variance of *Z* to know its distribution. Specifically, let 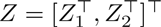, then it’s mean is

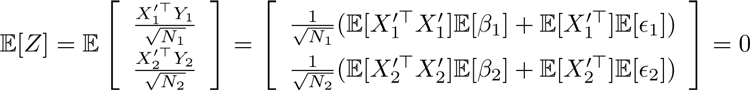

The within-population variance is:

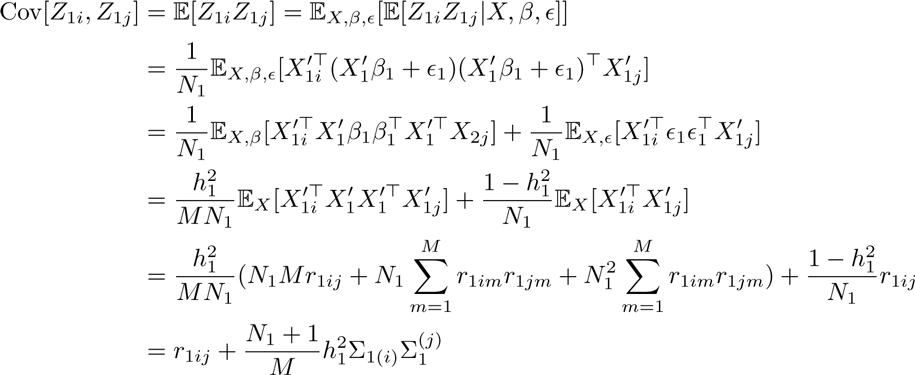

where 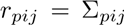 is the correlation coefficient of SNP *i* and *j* in population *p*. Similarly, the between-population variance is:

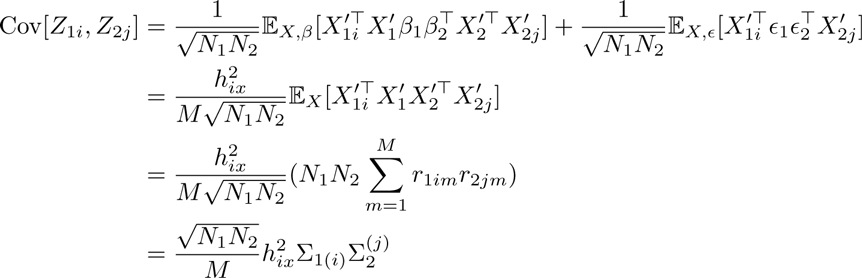

where 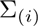 denotes the i’th row of *Σ* and 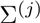 denotes the j’th column. The covariance of the *Z*-scores is thus

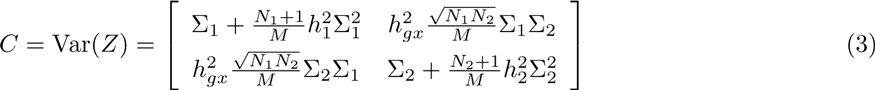

and 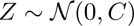.

### Genetic effect correlation

Let 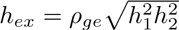. We modify the procedure above to use mean-centered instead of normalized genotype matrices and model the distribution of the effect sizes as

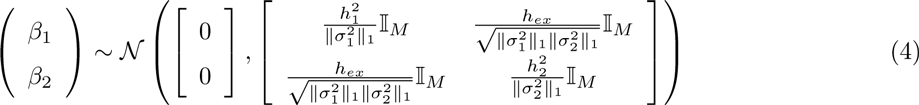

Notice that a linear model with efects sizes acting on un-normalized genotypes is the same as a linear model with effect sizes acting on normalized genotypes under the substitution 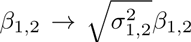. Therefore the covariance of *Z*-scores on the per allele scale can be immediately inferred from the prior derivation

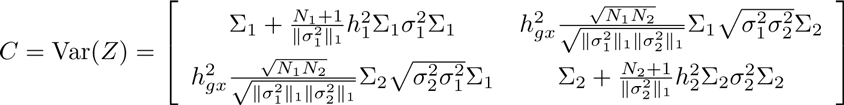

### Approximate maximum likelihood estimation

Let 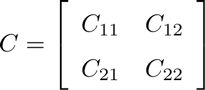 be either of the above covariance matrices written in block form. We approximately optimize the above likelihood as follows: first we find 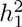 and 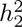 by maximizing the likelihood corresponding to *C_11_* and *C_22_*, then we find 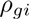 or 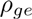 by maximizing the likelihood corresponding to *C_12_*:

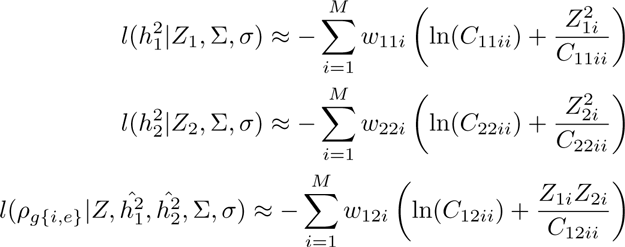

Because we are discarding between-SNP covariance information 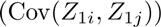, highly correlated SNPs will be overcounted in our approximate likelihood. As a simple example, notice that two SNPs in perfect LD will each contribute identical terms to the approximate likelihood, and therefore should be downweighted by a factor of 1/2. The extent to which SNP *i* is over-counted is exactly the i’th entry in it’s corresponding LD-matrix product. Therefore we let 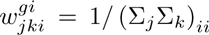 and 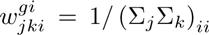 to reduce the variance in our estimates of the parameters 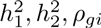 and 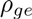.

Furthermore, rather than compute the full products 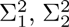 and 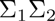 over all *M* SNPs in the genome, we choose a window size W and approximate the product by 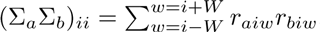. Though maximum likelihood estimation admits a straightforward estimate of the standard error via the fisher information, we found these estimates to be inaccurate in practice. Instead, we use block jackknife with block size equal to 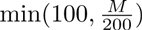 SNPs to ensure that blocks are large enough to remove residual correlations.

### Out of population prediction of phenotypic values

Consider using the results of a GWAS with perfect power in population 2 to predict the phenotypic values of a set of individuals from population 1. This defines the upper limit of the correlation of true and predicted phenotypic values. Let the true values of the effects sizes in population 2 be *β_2_*. Let the true phenotypes in population 1 be 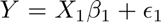 while the predicted phenotypes are 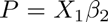. We are interested in the correlaiton of the predicted and true phenotypes 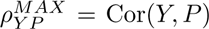. Notice that, given *X*, the true and predicted phenotype of each individual is an affine transformation of a multivariate normal random variable (*β*)

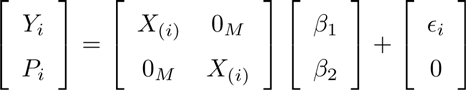

and therefore 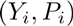 for individual *i* is multivariate normal with expected covariance matrix

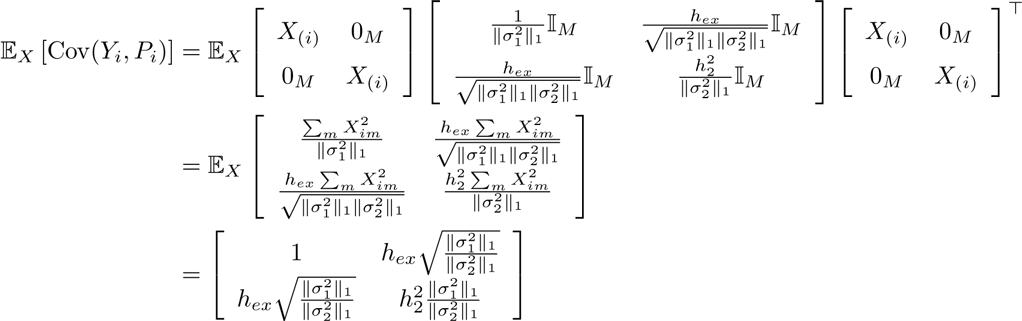

and therefore the expected correlation 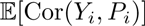 is 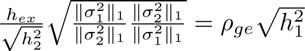. The expected population correlation tends to the sample correlation as the number of samples increases, therefore

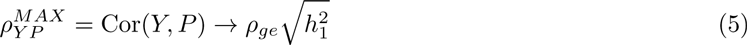

as 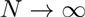

## Description of Supplemental data

Supplemental data include six figures and one table.

## Acknowledgements

The authors would like to acknowledge Lior Pachter and Hilary Finucane for insightful discussion about the problem. BCB is supported by the NSF GRFP. ALP is supported by NIH grant R01 HG006399. NZ is supported by NIH grant K25HL121295.

## Web Resources

*Popcorn* is available at https://github.com/brielin/popcorn

